# Conserved Cardiovascular Network: Bioinformatics Insights into Genes and Pathways for Establishing *Caenorhabditis elegans* as an Animal Model for Cardiovascular Diseases

**DOI:** 10.1101/2023.12.24.573256

**Authors:** Ashwini Kumar Ray, Anjali Priya, Md. Zubbair Malik, Thangavel Alphonse Thanaraj, Alok Kumar Singh, Payal Mago, Chirashree Ghosh, Shalimar, Ravi Tandon, Rupesh Chaturvedi

**Author notes:** Equal first author (Ashwini Kumar Ray, Anjali Priya). Corresponding to: (Dr. Ashwini Kumar Ray & Dr. Md. Zubbair Malik).

## Abstract

Cardiovascular disease (CVD) is a collective term for disorders of the heart and blood vessels. The molecular events and biochemical pathways associated with CVD are difficult to study in clinical settings on patients and *in vitro* conditions. Animal models play a pivotal and indispensable role in cardiovascular disease (CVD) research. *Caenorhabditis elegans*, a nematode species, has emerged as a prominent experimental organism widely utilised in various biomedical research fields. However, the specific number of CVD-related genes and pathways within the *C. elegans* genome remains undisclosed to date, limiting its in-depth utilisation for investigations. In the present study, we conducted a comprehensive analysis of genes and pathways related to CVD within the genomes of humans and *C. elegans* through a systematic bioinformatic approach.

A total of 1113 genes in *C. elegans* orthologous to the most significant CVD-related genes in humans were identified, and the GO terms and pathways were compared to study the pathways that are conserved between the two species. In order to infer the functions of CVD-related orthologous genes in *C. elegans, a* PPI network was constructed. Orthologous gene PPI network analysis results reveal the hubs and important KRs: *pmk-1, daf-21, gpb-1, crh-1, enpl-1, eef-1G, acdh-8, hif-1, pmk-2,* and*aha-1 in C. elegans.* Modules were identified for determining the role of the orthologous genes at various levels in the created network. We also identified 9 commonly enriched pathways between humans and *C. elegans* linked with CVDs that include autophagy (animal), the ErbB signalling pathway, the FoxO signalling pathway, the MAPK signalling pathway, ABC transporters, the biosynthesis of unsaturated fatty acids, fatty acid metabolism, glutathione metabolism, and metabolic pathways. This study provides the first systematic genomic approach to explore the CVD-associated genes and pathways that are present in *C. elegans,* supporting the use of *C. elegans* as a prominent animal model organism for cardiovascular diseases.

## 1. Introduction

Cardiovascular Disease (CVD) is a collective term for disorders of heart and blood vessels that include coronary heart disease, hypertension, and heart failure along with other heart and vascular system-related diseases such as atherosclerosis and deep venous thrombosis. CVDs are the leading cause of global mortality ^1^ with 17.9 million deaths each year ^2^which is expected to rise to 23.6 million by 2030^3^, according to the reports of WHO four out of five CVD patients die because of heart attack or myocardial infarction every year ^4^. CVD is influenced by a multitude of risk factors, prominent among these are lifestyle factors such as being overweight, having high blood sugar, increased blood pressure, smoking and unhealthy lifestyles along with genetic risk factors such as a family history of CVD that contribute to the development of CVD ^5^. A thorough understanding of the pathophysiology and pathological manifestations of CVD is required for developing efficient treatment strategies or preventative measures and lowering the rate of incidence and morbidity.

The molecular events and biochemical pathways associated with CVD are difficult to study in clinical settings on patients and *in vitro* conditions. *In vitro*, studies don’t give such complete information on the pathogenesis of organ system level ^6^. Thus, Numerous aspects of human cardiovascular disease (CVD) are not fully comprehended. Given that cellular metabolism is a complex network of biochemical pathways, gaining a detailed understanding of the genes associated with the disease within these pathways is essential for unraveling its complexities. A better understanding of interacting partners is a key aspect of translational research to develop a therapeutic strategy ^7^. Therefore, to better understand the molecular and biochemical pathways it is crucial to understand changes in the molecular events causing CVD by employing animal models.

Animal models play a very essential role in the research area of CVD. For example, Neuberger et al. studied chronic atrial dilation and atrial fibrillation using a goat model ^8^. Human atherosclerosis is studied using rabbits and pigs ^9,10^. Similarly, Canine, primate and rodent models have also been employed to study CVD ^11^. However, mammalian animal model systems have their limitations of high experimental cost, ethical issues, limited genomic tools available, complicated operation, and high reproductive environment demands despite having genomic relatedness ^12^.

The nematode *Caenorhabditis elegans* (*C. elegans*) ^13^ has found extensive application in diverse domains such as developmental and evolutionary biology, genetics, and intricate investigations into disease studies. ^14–17^. The genomic comparison of human and *C elegans* describes that the majority of human disease genes and pathways are found in *C. elegans* ^18^ . Approximately 40 - 75% of genes linked to human diseases show a certain level of similarity (E < 10 −10 in BLASTP searches) with genes identified in *C. elegans*^19–24^. A comprehensive proteomic analysis of 18,452 *C. elegans* protein sequences identified human gene homologs for approximately 83% of the *C. elegans* proteome ^25^. Moreover, *C. elegans* homologs were found to exist for ∼60–80% of human protein-coding genes ^23,25–27^. Moreover, in cases where a gene associated with a human disease does not have an orthologue in *C. elegans*, there is a significant likelihood that a homologous gene, a protein domain, or the elements of a related biological pathway—particularly if it involves a fundamental cellular function such as signal transduction, synaptic transmission, or membrane trafficking—are conserved. This conservation suggests that the *C. elegans* model system can provide valuable insights into human pathobiology^28,29^.

*C. elegans* is the first multicellular organism whose genome has been completely sequenced and thoroughly analysed ^30,31^. It possesses many lipid-binding proteins and transporters ^32^ along with biosynthesis of polyunsaturated fatty acids ^33^, ceramides ^34^ and phospholipids ^35–39^. The body wall muscle cells of *C. elegans* are useful for the study of human cardiomyocytes and their homologous structures and proteins ^40^. The nematode *C elegans* continues to be an excellent model for studying the organization, assembly and maintenance of the sarcomere ^40,41^. The major striated muscle of *C elegans* is found in the body wall and is required for the animal’s locomotion. Nevertheless, previous studies have proven that the close homology of proteins and structures of interest justify the study of nematode body wall muscle as a way to understand human heart muscle ^40^. It was pertinent, therefore, to understand the possibilities of using *C elegans* as a model for studying cardiovascular disease. However, how many CVD-related genes and pathways are present in the *C. elegans* genome is not known to date. Similarly, for its utilisation as an animal model for CVD, the genomic and functional conservation of pathways and genes from *C. elegans* to humans is unknown thus, limiting its in-depth utilization for investigations.

In the present study, we analysed the datasets related to human CVD to obtain DEGs which were used to identify essential key regulators by integrating the protein-protein interaction (PPI) network and their topological properties in humans. The key regulators and their first neighbour genes were used for the identification of orthologous genes present in *C. elegans.* The orthologous genes were to construct the PPI Network, key gene identification and module analysis, gene ontology and pathways analysis. Comparison of human and *C elegans* cardiovascular disease gene network, pathways and functional similarity could provide vital support to use *C elegans* as a model.

## 2. Methodology

A schematic representation of the methodology **(Figure 1)** adopted for obtaining differentially expressed genes related to CVD, identifying orthologous genes in *C. elegans,* comparing the pathways, developing the PPI network, topological analysis, and identification of novel key regulators are discussed in detail section-wise.

**Figure 1:**
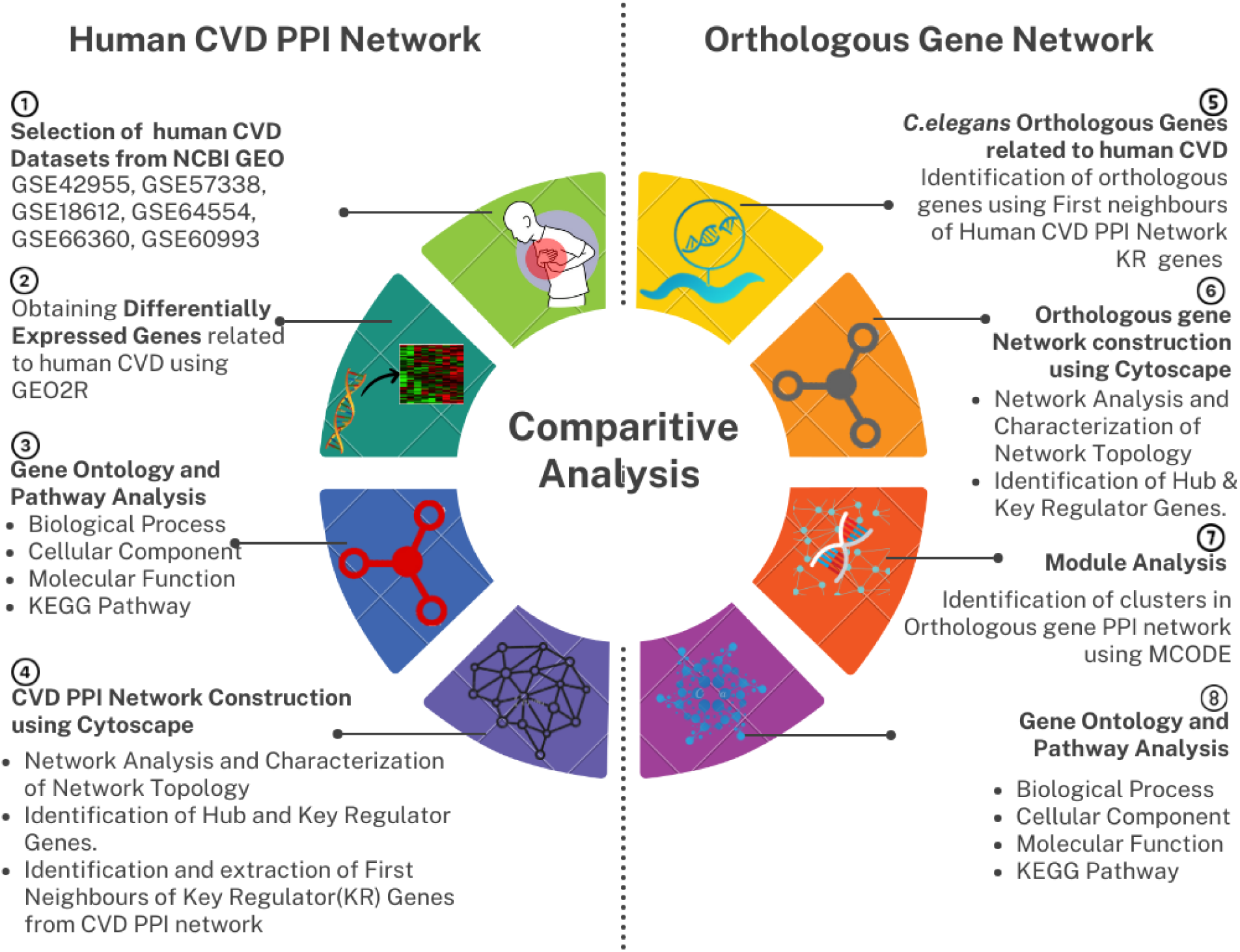
Schematic representation of the methodology of study showing the cardiovascular disease proteins were compared and analysed.

### 2.1 Selection of Cardiovascular Disease genes in human

There are several disorders that fall under the category of CVD. We selected the six most prevalent cardiac pathophysiologies for our analysis based on their significant contributions to CVD-related mortality which are as follows: cardiomyopathy, heart failure (HF), coronary artery disease (CAD), myocardial infarction (MI) and acute coronary syndrome (ACS) ^11,42^. For obtaining CVD-associated genes in humans, we studied the GEO microarray datasets from the National Centre for Biotechnology Information (NCBI). Six datasets GSE42955, GSE57338, GSE18612, GSE64554, GSE66360, and GSE60993 belonging to cardiomyopathy, HF, CAD, MI and ACS were selected for the study. Basic information on GEO Datasets used in the study is given in **Table 1**. The gene expression scores for the selected datasets were obtained by comparing data from patients with diseases and healthy individuals using the GEO2R tool which is an online analytical tool with an inbuilt R programme (https://www.ncbi.nlm.nih.gov/geo/geo2r/) ^43^. All the datasets were normalised and the statistical threshold of the p-value was set as ≤ 0.05. Lastly, Log fold change was used as the primary index for DEG screening. In this investigation, Log fold change |0.25-1.5| was used as a DEG screening condition. The obtained up and downregulated genes of CVD were further used for pathway analysis and construction of the PPI Network.

**Table 1:**
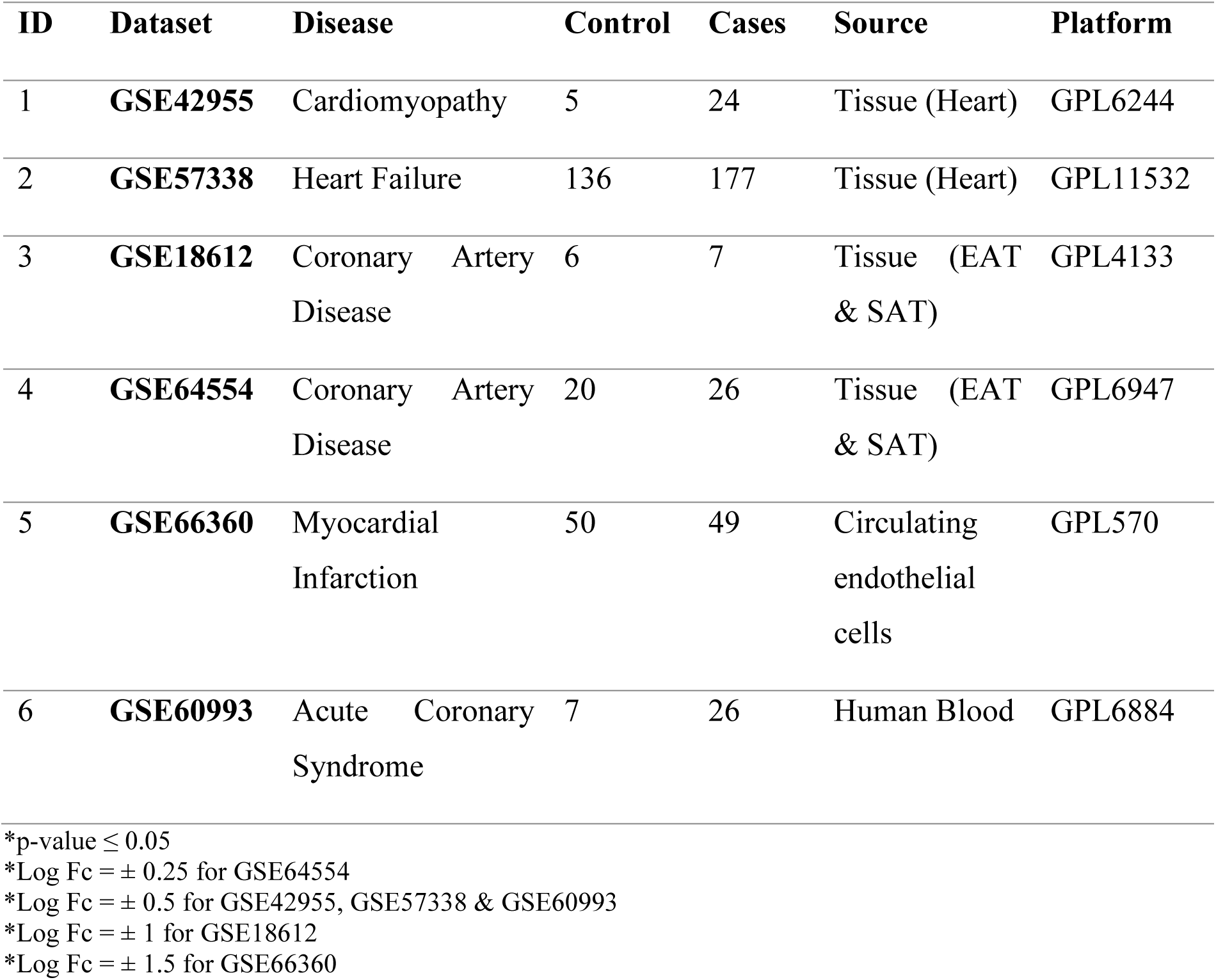
Basic information on GEO Datasets used in the study.

### 2.2 Human CVD PPI Network construction and Hub Gene identification

For understanding the regulatory function of the genes, the protein-protein interactions (PPI) network has been constructed. The human CVD PPI network was constructed using STRING database (https://string-db.org/) ^44^. The STRING database’s physical interaction data was then utilized for the construction, visualization, and analysis of networks using the Cytoscape 3.6.0 platform^45^. Finally, after eliminating isolated nodes and duplicate edges, a PPI network comprising 1099 upregulated and 815 downregulated genes was constructed in Cytoscape. The networks topological properties were analysed using Network Analyser ^46^ and Cytohubba ^47^ plugin of Cytoscape.

The network’s most impactful nodes are recognized as its hubs. Nodes with the highest degrees and correspondingly high centrality values were pinpointed as the hubs within the PPI network associated with CVD. To identify the key regulators (KRs) within the PPI network, the initial step involved identifying the network’s crucial genes based on their degrees (*k*) and centrality measures (*Cc, CB*). The top 10 genes of different topological properties were compared to identify KR genes48,49.

### 2.3 Identification of CVD related orthologous genes in *C. elegans*

In order to investigate how many potential CVD-related genes and pathways are present in the *C. elegans* genome, the orthologous genes of most significant CVD-associated genes were identified. This was achieved by identifying the first Neighbours of KRs of the human CVD PPI network and finding its orthologous in *C. elegans*. The first neighbour nodes of KR genes in the human interactome are defined as proteins directly and physically interacting with KR proteins ^50^. First neighbours play the same important roles in numerous PPI networks and signalling pathways as the associated KR proteins ^50^. Therefore, for identifying orthologous genes of most significant human CVD related genes, first neighbours of the essential key regulator genes were identified and extracted from the human CVD PPI Network. The identified 648 first neighbour genes that were highly significant were used for identification of orthologous genes present in *C. elegans*. g:Profiler that allows mapping of orthologous genes across species has been a popular tool for identification of *C. elegans*-human orthologs ^51^. The HGNC approved symbols of CVD genes of human is queried in g:Orth tool of g:Profiler (https://biit.cs.ut.ee/gprofiler/orth) to obtain the *C. elegans* genes (accessed on 15 December 2021).

### 2.4 Orthologous gene PPI Network construction and analysis

Animal models are employed to study the roles of specific genes or proteins associated with a disease with the speculation that the roles may be similar in humans. Comparison of human and model organism interactome can suggest that the proteins under study might play similar roles in human disease if their interaction is highly conserved ^52^. Therefore, we constructed the PPI network of CVD-related *C. elegans* orthologous genes. For this, we utilized the STRING database (https://string-db.org/) ^44^. First, an interactome network of orthologous genes was constructed on STRING. The physical interaction data of STRING was then used to for network construction, visualisation and analysis through the Cytoscape 3.6.0 platform ^45^. Finally, a PPI network of 581 nodes with 2519 connections (edges) was constructed on the Cytoscape after the removal of isolated nodes and duplicate edges.

Defining the fundamental topology of a network is generally done by analysing its topological parameters, especially *P*(*k*), *C*(*k*), and *CN* (*k*)^53^. So, first, the fundamental topology of the orthologous gene network was defined by analysing the above three parameters using the Network Analyzer app ^46^ in Cytoscape 3.6.0 taking the network as an undirected network. The three parameters *P*(*k*), *C*(*k*), and *CN* (*k*) of the network were analyzed by plotting against the degree *k* and their distribution behaviour was analysed to define the topologies of the orthologous gene network. The centrality measures which include Closeness centrality (*Cc*) and betweenness centrality (*CB*) of every node in the network were analysed, using the Network Analyzer app and CytoHubba app ^47^in Cytoscape 3.6.0 to study the influence and control capabilities of the nodes in signal processing and their dominance in network integration.

### 2.5 Identification of essential Key Regulators of orthologous gene network

To uncover key regulators within the orthologous gene PPI network, we initially identified the network’s crucial genes based on their degrees (*k*) and centrality measures (*Cc, CB*). Accordingly, nodes with the highest degrees also presented higher centrality values, thus identifying them as the most influential nodes and designating them as *hubs* within the network. Thus, key regulators with significant roles in the integration and stability of the PPI network would likely be found among these highly influential nodes and *hubs* of the network.

Hence, a group of top highest degree nodes, or hubs, that had the largest impact on the network from the level of the core network up to the level of motifs, were used to identify the network’s important regulators. The hubs functioning as the backbone of the system-level structure and represented at every hierarchical level were found by manual hub tracing and were regarded to be significant regulators.

### 2.6 Analysis of Modules

Modules within a large PPI network are characterized as a collection of statistically and functionally significant interacting genes. We constructed modules, using MCODE (Molecular Complex Detection) version 1.5.1 which follows the principle that highly connected regions (or clusters) of interaction networks are often complexes ^54^. MCODE tool is designed to detect clusters within a network that represent highly interconnected regions.. The default MCODE parameters i.e., “Degree cut-off = 2,” “node score cut-off = 0.2,” “k-score = 2,” and “max. depth = 100.” were used for network scoring and cluster finding. Significant modules were identified and filtered out based on MCODE score.

### 2.7 Comparative Pathway analysis of Human and *C. elegans*

Pathway-based analysis is critical for understanding the interaction of diseases at the molecular level. DAVID (https://david.ncifcrf.gov/) which is a comprehensive set of functional annotation tool was used to identify GO terms and Pathways ^55^. CVD related DEG in case of human and *C. elegans* orthologous genes were used to perform gene enrichment analysis. Comparison of pathways and GO terms of human CVD related genes and *C. elegans* orthologous genes was done to identify common pathways.

## 3. RESULTS

### 3.1 Identification and selection of CVD-related genes in humans

For gaining insights into the genes and pathways of CVD in model organism *C elegans* , six datasets belonging to cardiomyopathy, HF, CAD, MI and ACS were selected on the basis of high prevalence (Prevalence - CAD & ACS: 30%, MI: 5-30% for different age group, HF: 15-25% relative to age, cardiomyopathy: 0.2-0.5%) and its contribution to CVD-related mortality (Annual mortality rate – CAD & ACS: 5-6% per 1,00,000 population, MI: 34-42%, HF: 15%, Cardiomyopathy- 6-8%) ^11,42^. The details of all datasets used in the study are given in **Table 1**. The selected datasets were analysed using GEO2R. For the screening of DEGs, the statistical threshold of p-value was set as ≤ 0.05 and Log fold change was used as the primary index for filtering out the genes. Common DEGs shared by two datasets were identified and considered only once in the final gene list by removing duplicates. Finally, a total of 1099 upregulated and 815 downregulated genes associated with human CVD were retrieved by analysing the selected datasets **(supplementary file 1)**. The obtained DEGs were used for further study.

### 3.2 Human CVD-related gene network construction and *hub gene* identification

PPI network refers to the biochemical and biological activities of proteins in cells. It adds considerably to our understanding of cell physiology and disease association in the fields of biology and bioinformatics ^56^. Therefore, a PPI network was constructed using 1099 upregulated and 815 downregulated genes identified to be associated with human CVD. In the PPI network, proteins are denoted by nodes and the association between the proteins is given by undirected edges. The results of network analysis reveal that the CVD PPI network comprises 1626 interacting nodes and 16903 edges (**supplementary figure 1**).

Topological properties define the structural properties of a complex biological network ^57^. Therefore, in order to understand the essential behaviour of the CVD PPI network we studied the topological properties using the Network Analyzer app and CytoHubba app in Cytoscape. The “*hubs*” of the network are defined as the nodes with the highest degree in the system and tend to have an essential role in the PPI network. After analysis, we identified the top 10 most important *hub* genes (*IL6, IL1B, PTPRC, MYC, FN1, ITGAM, TLR4, STAT3, JUN* and *CXCL8)* from the CVD PPI network as per the decreasing value. Similarly, analysis of topological properties of the CVD PPI Network resulted in the top 10 centrality measures (Betweenness and Closeness centrality) as listed in **Table 2**.

**Table 2:**
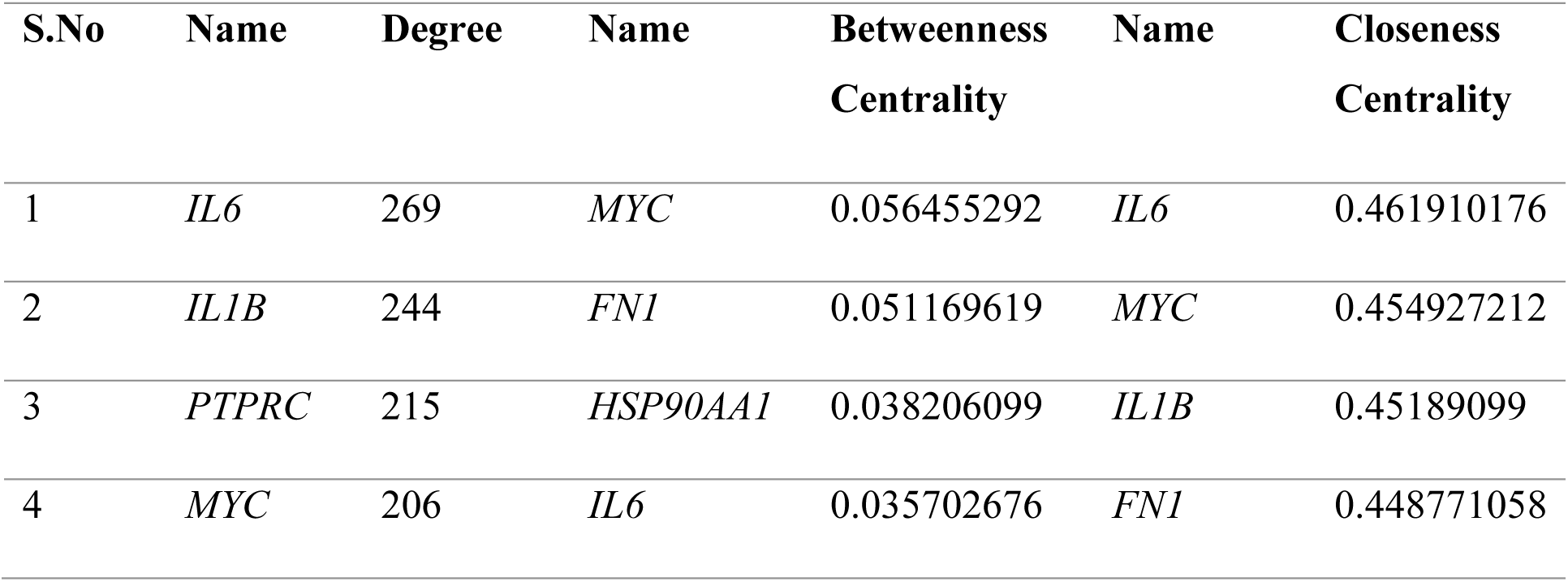
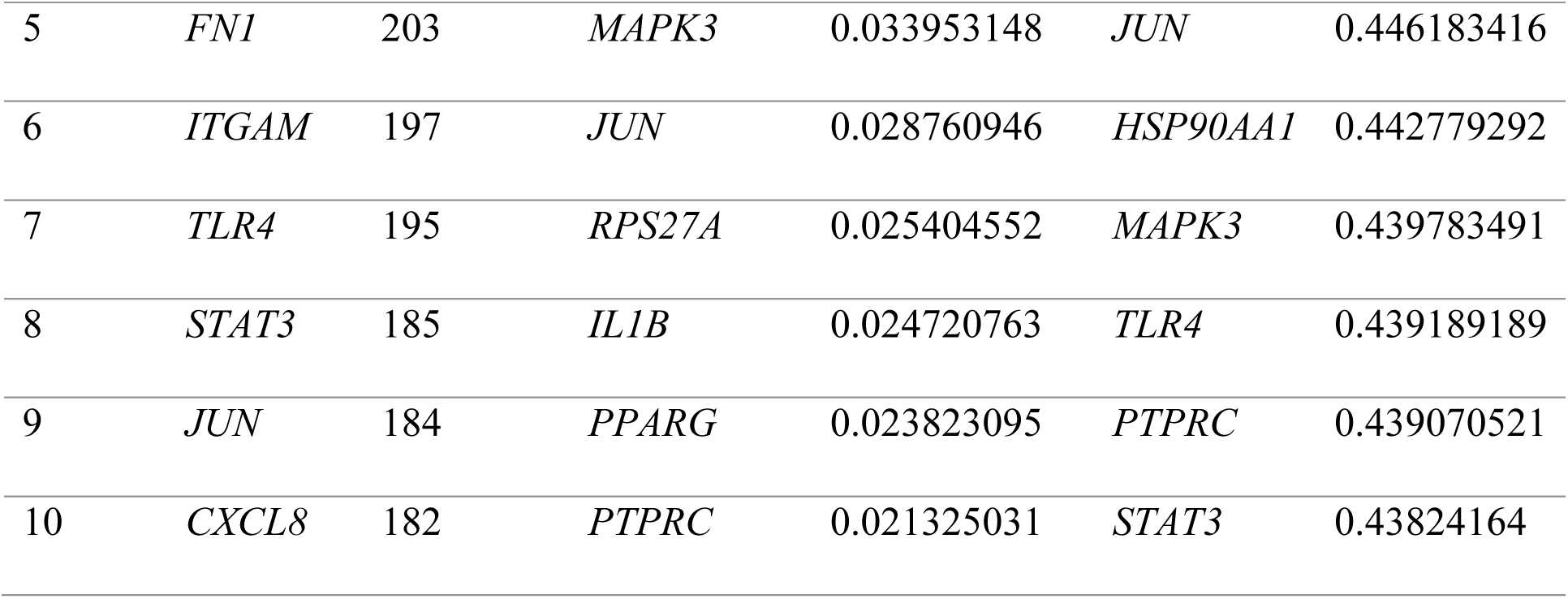
Top 10 highest Degree *(Hub genes),* Betweenness & Closeness centrality genes of the CVD PPI Network.

KR genes in a PPI network refer to a subset of genes that play pivotal roles in orchestrating and controlling various cellular processes through their interactions with other proteins. These genes exert significant influence over the network’s dynamics and functionality, often acting as central hubs or master regulators ^58^. Therefore, for the identification of essential KRs serving as the backbone of the CVD PPI network, the top 20 genes from three distinct topological features (degree, betweenness and closeness centrality) were compared **(Figure 2A, 2B, 2C)**. Eleven key regulators of the Human CVD that were common for all three attributes **(Figure 2D)** were identified as follows: *JUN, HSP90AA1, TLR4, CXCL8, MAPK3, PTPRC, IL6, IL1B, STAT3, MYC* and *FN1* indicating that these genes might play important regulatory functions in CVD development and progression in human. Seven out of eleven KRs were found to be upregulated with *IL1B* having maximum fold change (2.87) while four genes were downregulated with *HSP90AA1* having fold change of -1.21 **(Figure 2E)**. The expression of KRs in control and disease groups is presented by a box plot in **Figure 3**. The expression of *IL1B*, *JUN*, *FN1*, *CXCL8*, *TLR4*, *PTPRC* and *MAPK3* were significantly higher in diseased condition than in control while the expression of *HSP90AA1*, *IL6*, *STAT3* and *MYC* was found to be lower in diseased than control (P- value ≤ 0.05 for all the case).

**Figure 2:**
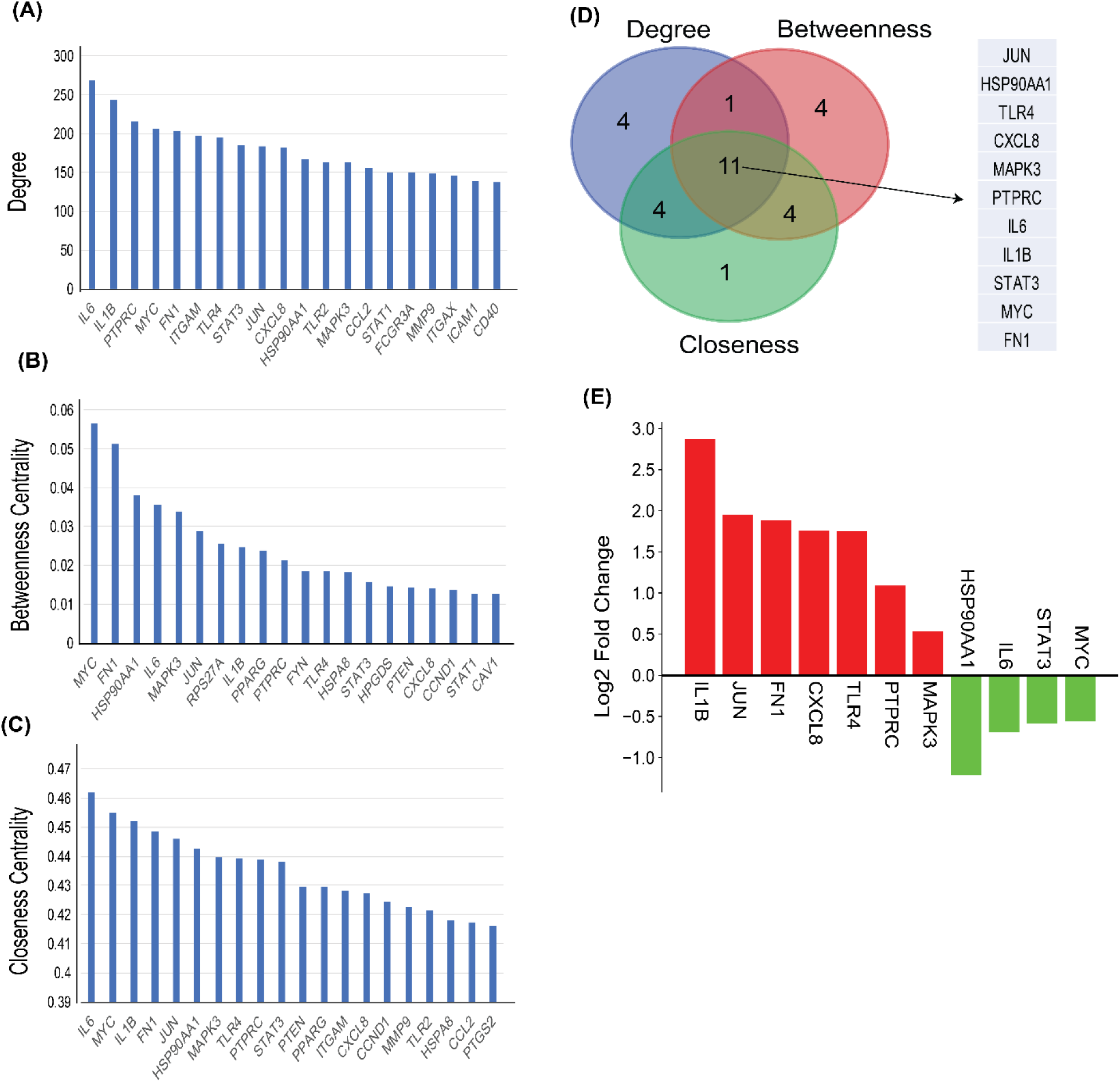
Identifying of key regulators using three attributes of CVD PPI Network. **(A-C)** Topological properties of the top 20 DEGs in terms of centrality measurements of degree, betweenness, and closeness. (**D**) Intersections among top ten genes having the highest centrality values of closeness, betweenness and degree. 11 common genes among the top 20 genes of each of topological parameter**. (E)** Bar plot showing the fold change for expression of 11 key regulators in individual healthy control and CVD patients. The red bar shows log2 foldchange of upregulated genes and green bars shows log2 foldchange of downregulated genes.

**Figure 3:**
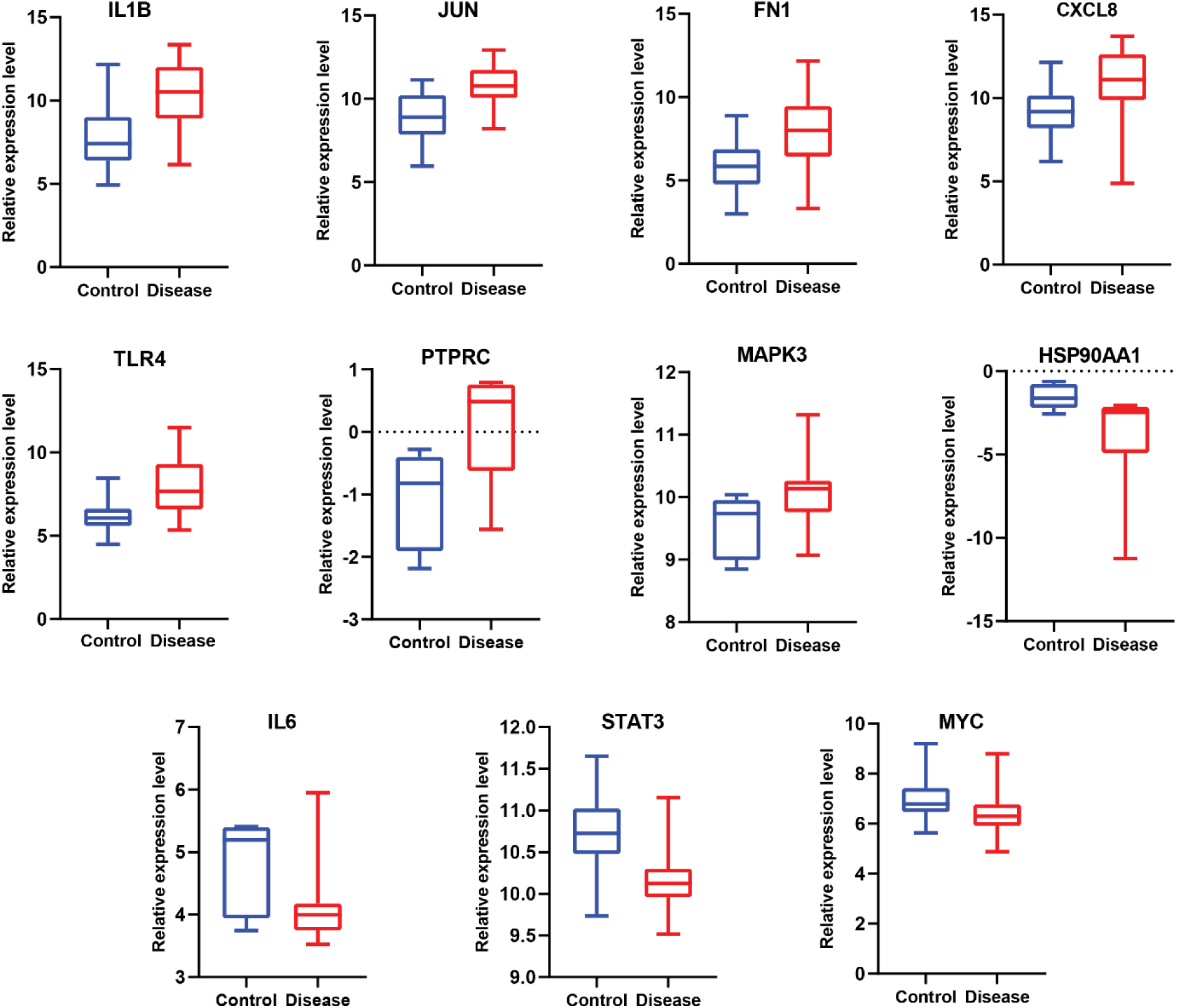
Relative **e**xpression of 11 key regulators normalized (norm) signal values between healthy controls and CVD patients. IL1B: interleukin 1 beta; JUN: Jun proto-oncogene, AP-1 transcription factor subunit; FN1: fibronectin 1; CXCL8: C-X-C motif chemokine ligand 8; TLR4: toll-like receptor 4; PTPRC: protein tyrosine phosphatase, receptor type C; MAPK3: mitogen-activated protein kinase 3; HSP90AA1: heat shock protein 90 alpha family class A member 1; IL6: interleukin 6; STAT3: signal transducer and activator of transcription 3; MYC: v-myc avian myelocytomatosis viral oncogene homolog.

### 3.3 Orthologous genes of human CVD in *C. elegans*

Completion of the *C. elegans* genome sequence in 1998 ^59^ demonstrated that roughly 38% of worm genes have a human ortholog ^60^. The percentage of human disease-related genes that have at least minor similarity (*E* < 10^−10^ on BLASTP searches) with *C. elegans* genes ranges between 40- 75% ^19–24^. For identifying orthologous genes of most significant human CVD-related genes, first neighbours of the essential key regulator genes were identified and extracted from the human CVD PPI Network. The first neighbour nodes of KR genes in the PPI network are the proteins that are directly and physically interacting with KR proteins ^50^. Therefore, we selected the first neighbour genes for *C. elegans* orthologue identification as they have similarly important roles in numerous PPI networks and signalling pathways as the associated KR proteins. A total of 637 first neighbour genes were found to be associated with the 11 KR proteins in the human CVD PPI network. Finally, 1113 *C. elegans* genes that were orthologous to 648 human CVD-related genes were identified **(supplementary file 2)**. The identified orthologous genes were used for construction PPI network and comparative pathway study.

### 3.4 Orthologous gene network construction and analysis

Many genes of different diseases in the genome of *C. elegans* have been extensively studied and their functions are well characterized. By studying the PPI network of CVD-related orthologous genes in *C. elegans*, we can infer the functions. Therefore, a PPI Network was constructed through STRING using the total *C. elegans* orthologous genes (**Supplementary Figure 2**). In the PPI network, proteins are represented by nodes, and the relationship between proteins is shown by undirected edges.

Further, to understand the structural properties of the PPI network based on network topology and connectivity patterns, Orthologous gene network analysis was done using Network Analyzer app and the Cytohubba plugin of Cytoscape. The “hubs” of the network are defined as the nodes with highest degree in the system. After analysis, we identified top 10 most important hub genes (*aha- 1, daf-21, pmk-1, gpb-1, pmk-2, cyb-2.1, cyb-2.2, hif-1, eef-1G* and *cyb-1)* from the PPI network. Similarly, analysis of topological properties of the CVD PPI Network resulted in the top 10 centrality measures (Betweenness and Closeness centrality) as listed in **Table 3**. **Figure 4A, 4B, 4C & 4D** shows the interaction of top 10 different Topological properties of CVD orthologous gene network.

**Figure 4:**
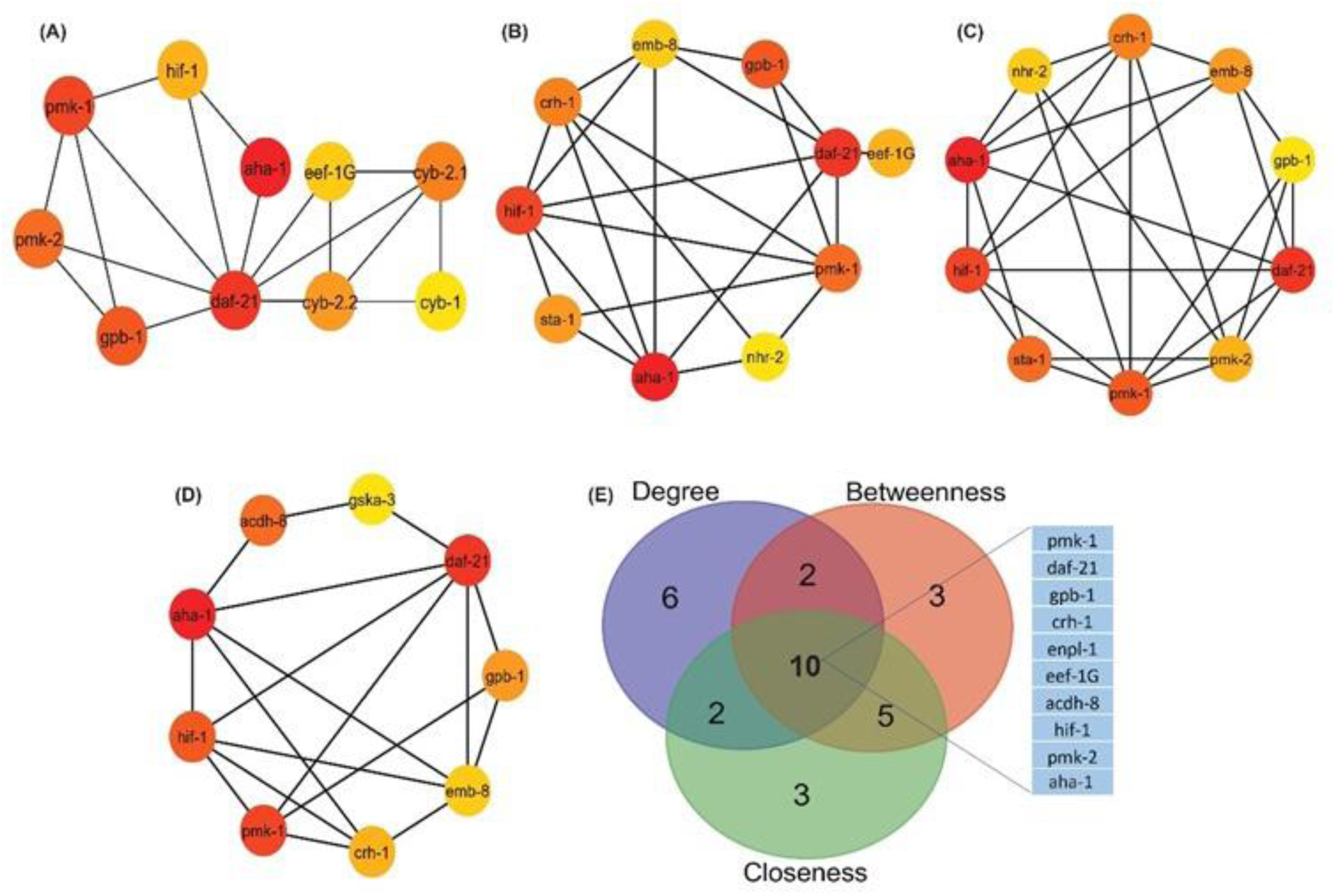
C. elegans orthologous gene PPI regulatory network of the top ten hub genes of CVD. Regulatory network of the top ten (**A**) Degree genes (**B**) Betweenness centrality genes **(C)** Closeness centrality genes **(D)** Bottleneck genes. (**E**) Intersections among top 20 genes having the highest degree and centrality values of closeness, betweenness. 10 common genes among the top ten genes of each of topological parameter

**Table 3:** Top 10 highest Degree *(Hub genes),* Betweenness & Closeness centrality genes of the *C. elegans* orthologous genes network.

| S.No | Name | Degree | Name | Betweenness Centrality | Name | Closeness Centrality |
| --- | --- | --- | --- | --- | --- | --- |
| 1 | *aha-1* | 166 | *aha-1* | 0.440878382 | *daf-21* | 0.442748092 |
| 2 | *daf-21* | 58 | *daf-21* | 0.148872762 | *aha-1* | 0.441400304 |
| 3 | *pmk-1* | 57 | *hif-1* | 0.081568412 | *hif-1* | 0.411639461 |
| 4 | *gpb-1* | 49 | *gpb-1* | 0.063156592 | *emb-8* | 0.400276052 |
| 5 | *pmk-2* | 48 | *pmk-1* | 0.060384451 | *sta-1* | 0.398078243 |
| 6 | *cyb-2.1* | 37 | *crh-1* | 0.05311299 | *pmk-1* | 0.394557823 |
| 7 | *cyb-2.2* | 36 | *sta-1* | 0.047005405 | *crh-1* | 0.392422192 |
| 8 | *hif-1* | 35 | *eef-1G* | 0.040100753 | *nhr-2* | 0.38410596 |
| 9 | *eef-1G* | 34 | *emb-8* | 0.037896232 | *pmk-2* | 0.379581152 |
| 10 | *cyb-1* | 32 | *nhr-2* | 0.036705488 | *enpl-1* | 0.366161616 |

For the identification of essential key regulators of the CVD orthologous gene PPI network, the top 20 genes from four distinct topological features (degree, bottleneck, betweenness and closeness centrality) were compared **(Figure 4E)**. Ten genes: *pmk-1, daf-21, gpb-1, crh-1, enpl-1, eef-1G, acdh-8, hif-1, pmk-2* and *aha-1* were found to be common for all the four attributes. When comparison of the identified key regulator nodes of human and *C. elegans* was done, it was found that two orthologous genes (*pmk-1* and *pmk-2*) of *MAPK3* which is an important key regulator of human CVD is serving as key regulators in *C. elegans* PPI network as well. *sta-1* which is orthologous to human gene *STAT3* is also among the top 10 topological properties of *C. elegans* PPI Network. All these results indicate that identified orthologous genes may play important role in elucidating the mechanisms of cardiovascular disease, which are currently poorly understood and *C. elegans* may serve as a good animal model.

### 3.5 Analysis of Modules in orthologous network

Modules represent groups of proteins that tend to interact with each other more frequently than with proteins outside the module ^61^. Modules can help us understand the dynamics of PPI networks. They may be transiently active in response to specific cellular conditions or environmental cues. Investigating module dynamics can reveal how the network adapts to changing circumstances. Hence, for determining the role of the orthologous genes in *C elegans* at various levels in the created network, the native or parent network was then divided into subnetworks or modules using the MCODE plug-in of the Cytoscape programme. The default MCODE parameters i.e., “Degree cut-off = 2,” “node score cut-off = 0.2,” “k-score = 2,” and “max. depth = 100.” were used for network scoring and cluster finding. A total of 25 modules were identified, out of which 9 modules having MCODE score ≥ 5 were considered significant and filtered from the PPI network. Interestingly, the identified key regulators were found to be part of these modules **(Figure 5).** Among the top 4 modules of Cardiovascular disease network, Module 1 has 24 nodes and 276 edges with a score of 24 **(Figure 5A)** ; Module 2 has 49 nodes and 361 edges with a score of 15.042 and contains four regulatory gene (*pmk-1,pmk-2,gpb-1,eef-1G*) **(Figure 5B)**; Module 3 has 17 nodes and 114 edges with a score of 14.250 and has one key regulator genes *acdh-8* **(Figure 5C)** and Module 4 has 8 nodes and 25 edges with a score of 7.143 **(Figure 5D)**.

**Figure 5:**
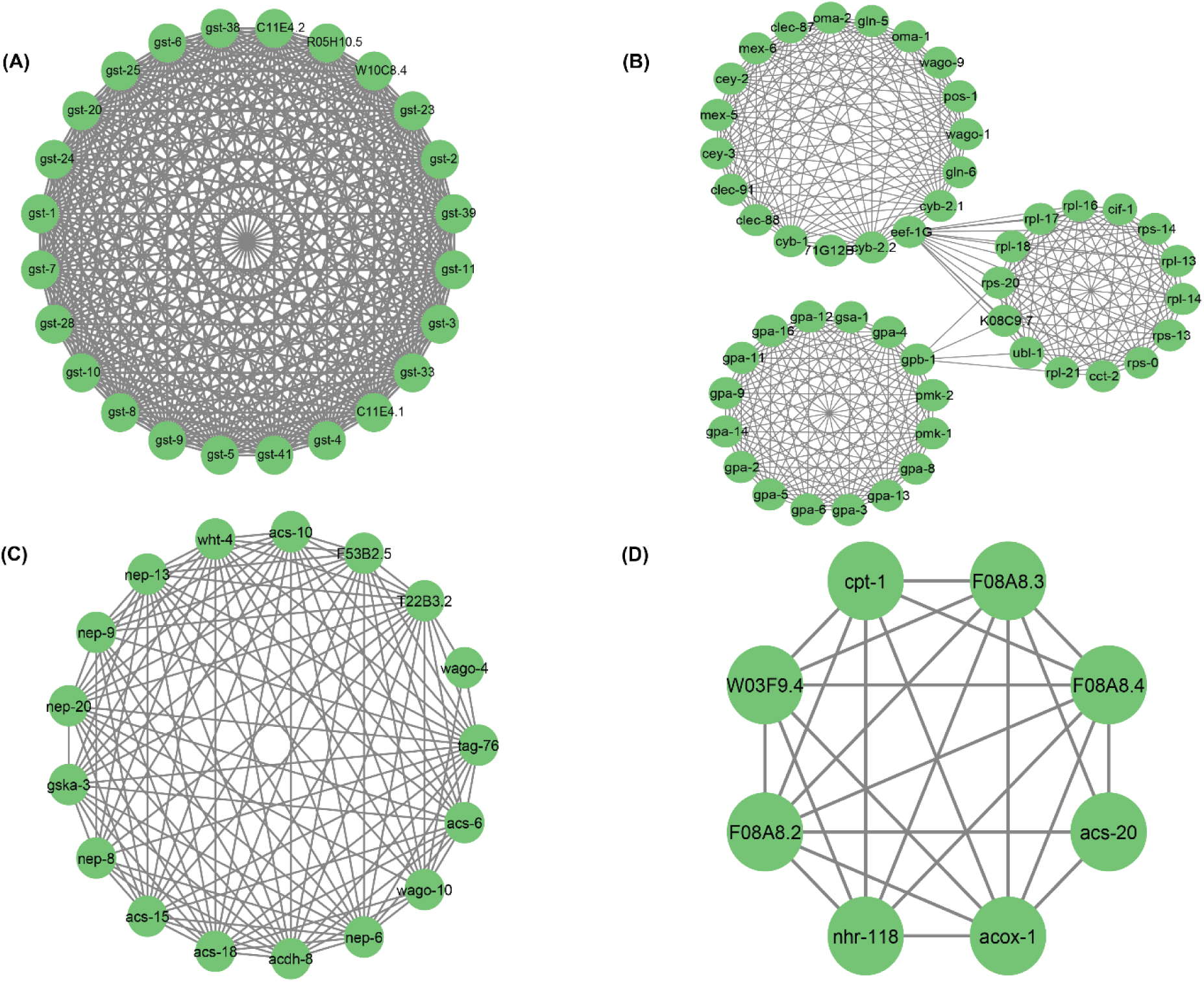
Top 4 modules of *C. elegans* orthologous gene PPI regulatory Network: **(A)** Module 1 has 24 nodes of genes and 276 edges as interaction with a score of 24.000; **(B)** Module 2 has 49 nodes and 361 edges with a score of 15.042; **(C)** Module 3 has 17 nodes and 114 edges with a score of 14.250 and **(D)** Module 4 has 8 nodes and 25 edges with a score of 7.143.

### 3.6 Comparative Pathway analysis of Human and *C. elegans*

In order to dissect the molecular mechanism of CVD in humans using *C. elegans* as an animal model, it is imperative to study the pathways that are conserved between the two species. In this study, we used the DAVID database to identify GO terms and KEGG pathways of the human CVD-related genes **(supplementary file 3 & 4)** and its orthologous genes in *C. elegans* **(supplementary file 5)**. A comparison of pathways and GO terms of humans and *C. elegans* was done to identify CVD-related pathways that are present in *C. elegans*. The pathway and GO analysis of 1099 upregulated and 815 downregulated human genes and 1113 *C. elegans* orthologous genes showed that they are highly enriched in similar pathways. There are 9 commonly enriched pathways between humans and *C. elegans* that include Autophagy – animal, ErbB signalling pathway, FoxO signalling pathway, MAPK signalling pathway, ABC transporters, Biosynthesis of unsaturated fatty acids, Fatty acid metabolism, Glutathione metabolism and Metabolic pathways. The list of pathways along with corresponding P-values in *C. elegans* and Humans are given in **Figure 6A**. The total number of CVD-related genes and orthologous genes that are involved in these pathways are given by bar plot in **Figure 6B**. The graph clearly demonstrates that CVD-related genes and their orthologous are involved in metabolic pathways.

**Figure 6:**
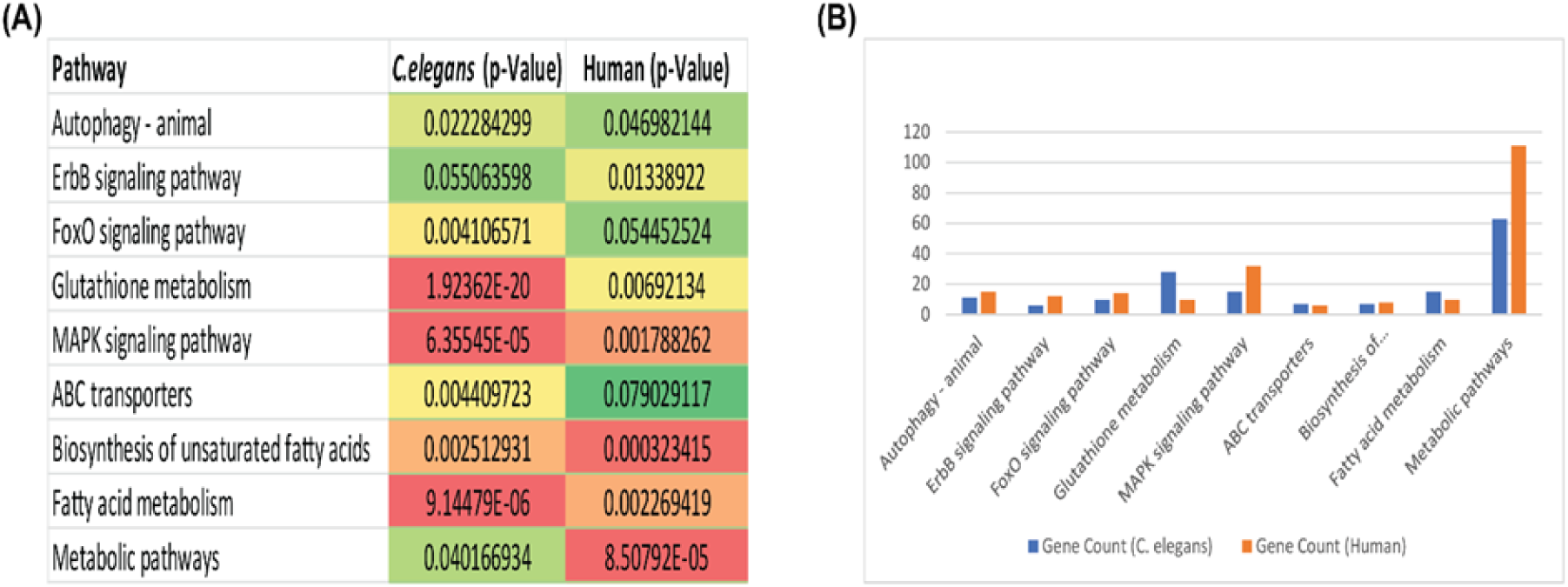
Functional comparison of the protein associated with CVD in humans and C. elegans. (**A**) Biological pathways of CVD proteins in humans and C. elegans using KEGG pathways with p-values. (**B**) Bar plot shows pathways comparison of the protein associated with CVD in humans and C. elegans. Y-axes represent no. of proteins and X- axis represents pathways.

Further, GO functional enrichment analysis also predicted similar functions in both the protein sets of human and *C elegans.* GO functions were classified into three categories. In each of these categories, several common functions were identified. We found 20 common Biological Processes describing the biological purpose of the gene or a gene product for carrying out a molecular function. Similarly, we also discovered 9 common Cellular Component and 13 common Molecular Functions through the comparative study of GO terms. Several important BP such as MAPK cascade, carnitine metabolic process, cell differentiation, chaperone-mediated protein folding requiring cofactor, fatty acid metabolic pathway, glutathione metabolic process etc. are common among *C. elegans* and humans. The bubble plot in **Figure 7** illustrates the GO terms and KEGG pathways for *C. elegans* orthologous genes. The red colour highlights the enrichment of similar GO terms and Pathways in humans and *C. elegans.* As a result of common pathways and GO terms present between humans and *C. elegans* we speculate that C. elegans could serve as an animal model for studying cardiovascular diseases.

**Figure 7:**
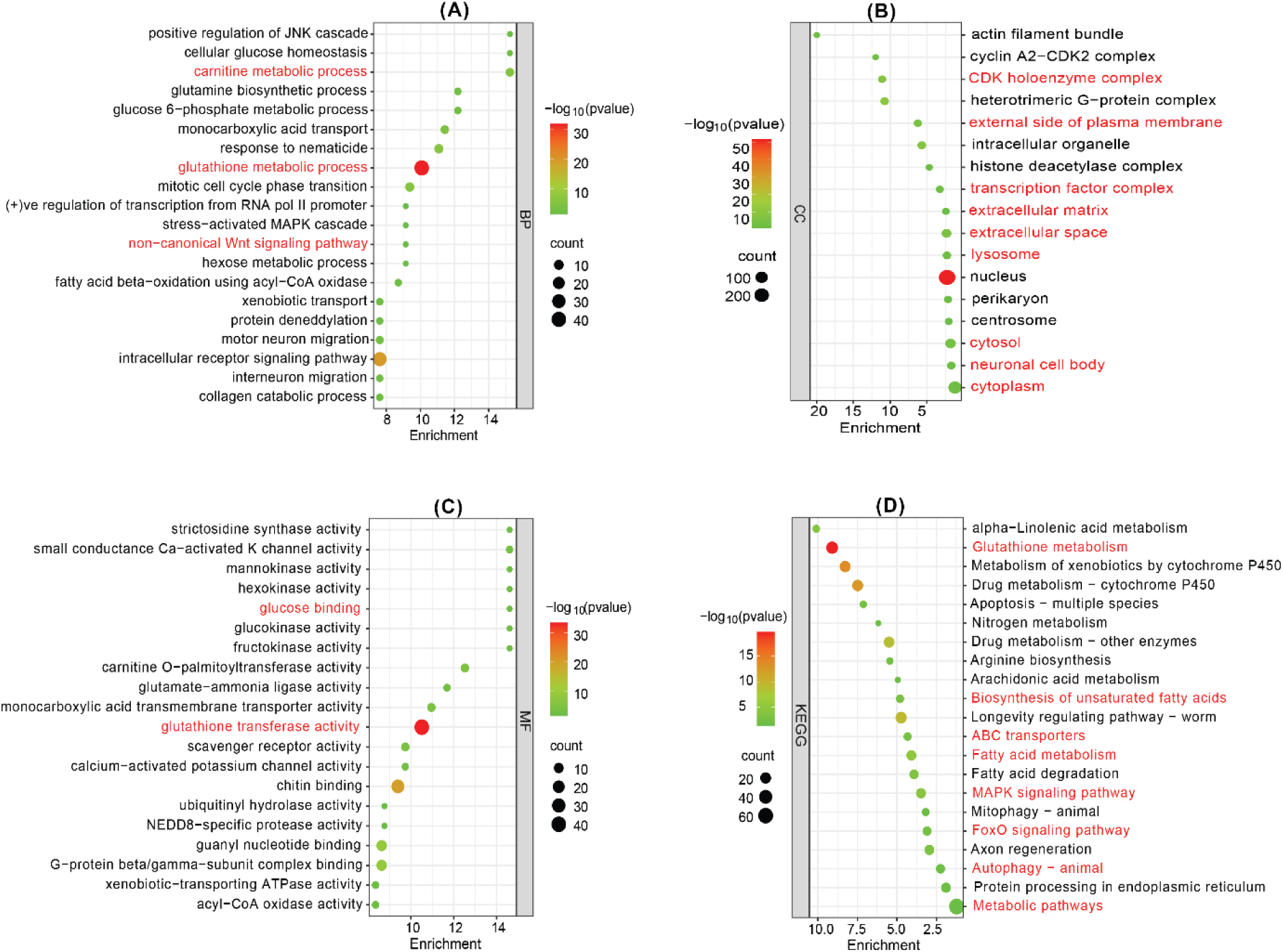
Functional enrichment analysis of *C. elegans* orthologous genes. **(A)** The significant enriched biological process of targeted *C. elegans* orthologous genes. **(B)** The significantly enriched cellular component, **(C)** The significant enriched molecular function and **(D)** KEGG Pathway. Dot size indicate count. The count represents the number of genes associated with each process. The dot color denotes the p values of process, and the x-axis represents the fold enrichment score.

## 4. DISCUSSION

With the rise in global incidences of CVD, it is becoming important to understand the CVD-related genes and pathways for gaining insights into the molecular and biochemical mechanism of the disease in order to develop efficient treatment strategies or preventative measures and lower the rate of incidence and morbidity. Various mammalian animal models are often employed to study CVD. In spite of having genomic relatedness, mammalian animal model systems have their own limitation such as high experimental cost, ethical issues, limited genomic tools available, complicated operation, and high reproductive environment demands ^12^. A simple well – known model organism such as *C. elegans* whose genome has been completely sequenced and thoroughly studied ^59^ can give a comprehensive genomic evaluation of a wide range of human diseases including CVD ^40,62–64^. However, to date, we lack information regarding the exact number of CVD-related genes and pathways within the *C. elegans* genome. Likewise, the extent of genomic and functional similarities between *C. elegans* and humans in relation to CVD remains unknown, which restricts its comprehensive use as an animal model for cardiovascular disease research.

Since it is crucial to recognize the potential of the model organism as a valuable candidate for studying CVD, we have used a bioinformatics approach for comparing the genes and pathways related to CVD of humans with those in *C. elegans*. We analysed microarray datasets of patients to obtain CVD-related genes. A total of 1099 upregulated and 815 downregulated human CVD-associated genes were identified from gene expression analysis. In order to understand the cell physiology and disease connection of these DEGs, we constructed a PPI network. The network derived from DEG exhibits a hierarchical structure, indicating a systematic organization with interconnected modules at various levels. Due to the hierarchical nature of the network, the synchronization among its components demonstrates critical functional regulations within the network. The network’s crucial regulators, denoted as significant genes or leading hubs, play a pivotal role in influencing motifs and governing module regulation, underscoring their profound biological significance. The leading hubs play a pivotal role with significantly important functions. They bring together lower degree nodes to effectively organize and regulate activities, fostering inter and intra-cross-talk among essential genes. Additionally, they contribute to sustaining network properties and stability, all while optimizing the processing of signals within the network.65,66.

After comprehensive network analysis using the Network Analyzer app and Cytohubba plugin of Cytoscape, we identified the eleven proteins as central or key regulators within the human CVD network which are as follows: *JUN, HSP90AA1, TLR4, CXCL8, MAPK3, PTPRC, IL6, IL1B, STAT3, MYC* and *FN1*. Our results demonstrate that the expression of *IL1B*, *JUN*, *FN1*, *CXCL8*, *TLR4*, *PTPRC* and *MAPK3* were significantly higher in the diseased condition than control while the expression of *HSP90AA1*, *IL6*, *STAT3* and *MYC* was found to be lower in diseased than control **(Figure 3).** There is increasing evidence suggesting that inflammation plays a role in the development of CVD. *IL1B* is a pro-inflammatory cytokine ^67^. Many research investigations have shown that individuals with heart failure display elevated levels of inflammatory cytokines in their bloodstream, including *IL-1B* ^67^. The product of the *CXCL8* gene (commonly known as IL8) belongs to the CXC chemokine family and serves as a primary mediator of the inflammatory response ^68^. Several research investigations have detected the presence of *IL-8* at locations of vascular damage, while separate studies have suggested that *IL-8* may be involved in different phases of atherosclerosis ^68^. In the findings by Rus et al. elevated concentrations of *IL-8* were reported within the atherosclerotic wall of human arteries ^69^. In another study by Liu et al., it was identified that foam cells originating from human atherosclerotic tissues had elevated concentrations of *IL-8* ^70^. Toll-like receptors (*TLRs*) are present in a variety of heart cell types, including cardiac myocytes, smooth muscle cells, and endothelial cells ^71^. They have been reported to be involved in a variety of functions in diseased conditions such as CVD, apoptosis, allergic diseases etc^72^. Several research investigations have provided evidence that *TLR4* activation leads to the upregulation of multiple genes associated with pro-inflammatory cytokines ^73,74^, which have significant roles in promoting myocardial inflammation, especially in conditions such as myocarditis^75^ , myocardial infarction, ischemia-reperfusion injury ^76^, and heart failure ^77^. Our result shows the upregulation of these inflammation-related genes except IL6 which is downregulated. The fact that these play a significant role as key regulators emphasizes the critical role of inflammation in CVD is an important finding of our study.

In addition to the identification of inflammation-related genes in the study, we have also uncovered non-inflammatory genes that bear significance in the context of CVD such as *JUN* is found to be a key player implicated in the prevention of stress-induced maladaptive remodelling of the heart^78^. Additionally, *MAPK3*, a participant in intracellular mitogen-activated protein kinase (*MAPK*) signalling cascades, is believed to play crucial role in the pathogenesis of cardiac and vascular diseases ^79^. *HSP90AA1*, an isoform of heat shock protein 90, has been associated with its role in cardiac remodelling ^80^. The presence of *PTPRC* has been noted in atherosclerotic plaque development ^81^, while *MYC* has exhibited involvement in cardiac myotropy and angiogenesis ^82^. *FN1* has been linked to coronary artery disease ^83^, and *STAT3* has been recognized for its cardioprotective functions ^84^. These genes, integral to cellular physiology, contribute significantly to diverse aspects of cardiovascular health and disease. Within the PPI network, key regulators (KRs) identified in this study offer valuable insights into the molecular mechanisms underpinning cardiovascular conditions. Consequently, these proteins emerge as promising targets for future research and therapeutic interventions in the development and progression of CVD. It is imperative to underscore the necessity for further experimental studies to validate these findings and elucidate the precise roles of these key regulators in cardiovascular health and disease.

For a better understanding of cell physiology and disease connection of orthologous genes in *C. elegans* we constructed the PPI network of orthologous genes. In the PPI network, proteins are represented by nodes, and the relationship between proteins is shown by undirected edges. Orthologous gene network analysis results identified the top 10 most important hub genes (*aha-1, daf-21, pmk-1, gpb-1, pmk-2, cyb-2.1, cyb-2.2, hif-1, eef-1G* and *cyb-1)* from the PPI network. Ten essential key regulator genes which are as follows: *pmk-1, daf-21, gpb-1, crh-1, enpl-1, eef-1G, acdh-8, hif-1, pmk-2* and *aha-1* were identified from the PPI network suggesting its crucial role. In diseased conditions such as cancer and various heart and lung diseases, cells and tissue suffer from low oxygen conditions (pathological hypoxia) ^85^. Studies have reported that hypoxia activates *hif-1* gene in *C. elegans* ^86,87^. Similarly, in case of humans, *HIF* (ortholog of *hif-1*) is activated in hypoxic environment. This activation plays a central role in tissue repair, ischemia and cancer ^88^. Additionally, results also show that ortholog of important CVD-related genes such as *MAPK3, HSP90AA1, HSP90AB1, HSP90B1* (*pmk-1, pmk-2, daf-21, enpl-1)* playing important role in ortholog gene network. HSP90 has been found to be associated to several CVD. Its client proteins have been found to be involved in cardiac disease pathways such as MAPK signalling, TNF-α signalling, etc ^89^. A study of these proteins and signalling pathways using *C. elegans* ortholog can provide better insights into the molecular mechanism associated with the disease.

Results of comparative pathway analysis showed that there are 9 commonly enriched pathways that are present between humans and *C. elegans*. Among the common pathways, two belonged to lipid metabolic pathways “Biosynthesis of unsaturated fatty acid” and “Fatty acid metabolism”. Pathways related to lipid metabolism play a very crucial role in CVD as dysregulation in any step can cause disorders such as diabetes, obesity, atherosclerosis etc. A study by Zhang et al. clearly establishes *C. elegans* as a promising model for the study of lipid metabolism and lipid metabolic diseases ^90^ which is consistent with our result as dysregulation in lipid metabolism pathways is a risk factor for CVD. *C. elegans* and humans also have Autophagy – animal, ErbB signalling pathway, FoxO signalling pathway, MAPK signalling pathway, ABC transporters, Glutathione metabolism and Metabolic pathways that are commonly enriched, exhibiting the potential to contribute in human CVD development and progression thus making *C. elegans* as a suitable model organism to study the molecular events and pathways related to CVD.

Autophagy can contribute to CVD by engaging in various crucial signaling pathways, such as PI3K/Akt/mTOR, IGF/EGF, AMPK/mTOR, MAPKs, p53, Nrf2/p62, Wnt/β-catenin, and NF-κB pathways ^91^. Recent investigations have uncovered the intricate mechanisms that underlie the therapeutic impact of NRG-1/ErbB signaling in addressing heart failure. By activating upstream signaling molecules like phosphoinositide 3-kinase, mitogen-activated protein kinase, and focal adhesion kinase, the activation of the NRG-1/ErbB pathway leads to elevated cMLCK expression and improved intracellular calcium cycling. The former serves as a regulator of the contractile machinery, while the latter initiates cell contraction and relaxation ^92^ .

FoxOs exhibit diverse functions in mitigating stress, determining cell fate, and regulating energy availability. Given the heart’s ongoing adjustments to diverse stresses and metabolic states, FoxOs emerge as pivotal players in cardiac physiology ^93^. The investigation into the pathogenesis of atherosclerotic vascular diseases involves exploring the roles of extracellular signal-regulated kinase (ERK), C-jun N-terminal kinase (JNK), and p38 MAPK in cardiac hypertrophy, cardiac remodeling post-myocardial infarction, atherosclerosis, and vascular restenosis ^94^. Additionally, the potential significance of Adenosine triphosphate (ATP)-binding cassette (ABC) transporters is underscored, given their involvement in crucial aspects such as cholesterol homeostasis, blood pressure regulation, endothelial function, vascular inflammation, and the processes of platelet production and aggregation ^95^. During the progression of many cardiovascular pathologies, there is a development of oxidative stress, marked by the generation of reactive oxygen species (ROS) and reactive nitrogen species (RNS). Reduced glutathione (GSH), a tripeptide present in all tissues, plays an active role in mitigating the oxidative impact of reactive species. The synthesis and/or regeneration of GSH are crucial for effectively responding to the heightened presence of oxidizing agents therefore, preventing or mitigating harmful reactive oxygen species (ROS) effects in CVDs ^96^

Further, GO functional enrichment analysis also predicted similar functions in both the protein sets of human and *C elegans.* In each of the GO categories, several common functions were identified. We found 20 common Biological Processes describing the biological purpose of the gene or a gene product for carrying out a molecular function. Similarly, we also discovered 9 common Cellular Component and 13 common Molecular Functions through the comparative study of GO terms. As a result of common pathways and GO terms present between humans and *C. elegans* that are related to CVD All these results indicate that *C. elegans* can be a potential animal model to study various pathways and genes that are related to CVD. Consequently, the CVD-related genes present in *C. elegans* should be considered as a starting point and further experimental investigations are required to fully explore *C. elegans* as a model organism for CVD.

## 5. CONCLUSION

In summary, we explored the applicability of *C. elegans* as an animal model to study CVD. We identified 1113 orthologous genes in *C. elegans* that are related to CVD in humans. We also explored the pathways and GO terms related to CVD that are common between humans and *C. elegans.* This study provides the first genomic view on CVD-related genes and pathways that are present in *C. elegans*, supporting its use as a prominent animal model for the study of CVD.

### Data availability

The datasets analysed during the current study are available in the GEO repository. It is a public free repository database, which stores a large number of gene functions and expressions. The working link are as follows:

GSE42955:https://www.ncbi.nlm.nih.gov/geo/query/acc.cgi?acc=GSE42955,

GSE57338:https://www.ncbi.nlm.nih.gov/geo/query/acc.cgi?acc=GSE57338,

GSE18612:https://www.ncbi.nlm.nih.gov/geo/query/acc.cgi?acc=GSE18612,

GSE64554:https://www.ncbi.nlm.nih.gov/geo/query/acc.cgi?acc=GSE64554,

GSE66360:https://www.ncbi.nlm.nih.gov/geo/query/acc.cgi?acc=GSE66360,

GSE60993:https://www.ncbi.nlm.nih.gov/geo/query/acc.cgi?acc=GSE60993

### Author contributions

AKR and MZM study concept and design, acquisition of data, analysis and interpretation of data, drafting of the manuscript, and statistical analysis. AP and MZM acquisition of data, analysis and drafting of the manuscript, statistical analysis, AKR, MZM, TAT, interpretation of data and drafting manuscript, statistical analysis, Shalimar, CG, RC, AKS, RT and PM, formulation of draft and critical revision of the manuscript for intellectual content. All authors contributed to the article and approved the submitted version.

## Acknowledgement

**Figure 1** was created using Canva

## Funding

A.K.R is supported by GIA Scheme of Department of Health Research (DHR), Indian Council of Medical Research (ICMR), Govt. of India (F. No. R.11013/12/2023-GIA/HR), National Heart, Lung, and Blood Institute (NHLBI) and Fogarty International Centre (FIC), NIH, USA grant D43TW009345 and IoE, 2022, FRP, University of Delhi.

## Competing interests

The authors declare no competing interests.

## Notes

https://www.ncbi.nlm.nih.gov/geo/query/acc.cgi?acc=GSE42955

https://www.ncbi.nlm.nih.gov/geo/query/acc.cgi?acc=GSE57338

https://www.ncbi.nlm.nih.gov/geo/query/acc.cgi?acc=GSE18612

https://www.ncbi.nlm.nih.gov/geo/query/acc.cgi?acc=GSE64554

https://www.ncbi.nlm.nih.gov/geo/query/acc.cgi?acc=GSE66360

https://www.ncbi.nlm.nih.gov/geo/query/acc.cgi?acc=GSE60993

### Competing Interest Statement

The authors have declared no competing interest.

### Summary of Updates

Authors affliation order is updated; Figure 2 revised.

